# Store-operated Ca^2+^ entry is involved in endothelium-to-mesenchymal transition in lung vascular endothelial cells

**DOI:** 10.1101/2024.12.06.627034

**Authors:** Aleksandra Babicheva, Ibrahim Elmadbouh, Shanshan Song, Michael Thompson, Ryan Powers, Pritesh P. Jain, Amin Izadi, Jiyuan Chen, Lauren Yung, Sophia Parmisano, Cole Paquin, Wei-Ting Wang, Yuqin Chen, Ting Wang, Mona Alotaibi, John Y.-J. Shyy, Patricia A. Thistlethwaite, Jian Wang, Ayako Makino, Y.S. Prakash, Christina M. Pabelick, Jason X.-J. Yuan

## Abstract

Endothelial-to-mesenchymal transition (EndMT) is a biological process that converts endothelial cells to mesenchymal cells with increased proliferative and migrative abilities. EndMT has been implicated in the development of pulmonary vascular remodeling in pulmonary arterial hypertension (PAH), a fatal and progressive lung vascular disease. Transforming growth factor β_1_ (TGF-β_1_), an inflammatory cytokine, is known to induce EndMT in many types of endothelial cells including lung vascular endothelial cells (LVEC). An increase in cytosolic free Ca^2+^ concentration ([Ca^2+^]_cyt_) is a major stimulus for cellular proliferation and phenotypic transition, but it is unknown whether Ca^2+^ signaling is involved in EndMT. In this study we tested the hypothesis that TGF-β_1_-induced EndMT in human LVEC is Ca^2+^-dependent. Treatment of LVEC with TGF-β_1_ for 5-7 days resulted in increase in SNAI1/2 expression, induction of EndMT, upregulation of STIM/Orai1 and enhancement of store-operated Ca^2+^ entry (SOCE). Removal (or chelation) of extracellular or intracellular Ca^2+^ with EGTA or BAPTA-AM respectively abolished EndMT in response to TGF-β_1_. Moreover, EGTA diminished TGF-β_1_-induced increase in SNAI in a dose-dependent manner. Knockdown of either STIM1 or Orai1 was sufficient to prevent TGF-β-mediated increase in SNAI1/2 and EndMT, but did not rescue the continuous adherent junctions. Blockade of Orai1 channels by AnCoA4 inhibited TGF-β-mediated EndMT and restored PECAM1-positive continuous adherent junctions. In conclusion, intracellular Ca^2+^ signaling plays a critical role in TGF-β-associated EndMT through enhanced SOCE and STIM1-Orai1 interaction. Thus, targeting Ca^2+^ signaling pathways regulating EndMT may be a novel therapeutic approach to treat PAH and other forms of pre-capillary pulmonary hypertension.

**New & Noteworthy:** EndMT has been reported to contribute to the pathogenesis of PH. In this study we aimed to determine the role of Ca^2+^ signaling in the development of EndMT in human lung vascular endothelial cells. Our data suggest that TGF-β_1_ requires store-operated Ca^2+^ entry through STIM1/Orai channels to induce SNAI-mediated EndMT. For the first time we demonstrated that TGF-β_1_-induced EndMT is Ca^2+^-dependent event while inhibition of STIM1/Orai interaction attenuated EndMT in response to TGF-β_1_.

## Introduction

Endothelial-to-mesenchymal transition (EndMT) is a phenotypical conversion of fully differentiated and functional endothelial cells (EC) to highly proliferative mesenchymal cells (e.g., myofibroblasts and smooth muscle-like cells). It has been reported that EndMT is involved in the development of pulmonary arterial hypertension (PAH)^1-4^. PAH is a devastating disease in which pulmonary vascular resistance (PVR) is increased by sustained pulmonary vasoconstriction, concentric pulmonary vascular remodeling, in situ thrombosis and obliterative intimal lesions. Pulmonary vascular remodeling is one of the main causes for the increased PVR and pulmonary arterial pressure (PAP). Concentric PA medial thickening is developed mainly due to the enhanced PA smooth muscle cell (SMC) proliferation, while lung vascular endothelial cells (LVEC) undergoing EndMT could be a source for highly proliferative and migratory mesenchymal cells contributing to the development of occlusive intimal lesions. We and other investigators established the presence of EndMT during the development and progression of pulmonary vascular remodeling in patients with PAH and animals with severe experimental pulmonary hypertension (PH)^2,5,6^.

Despite the fact that EndMT is widely studied, the molecular mechanisms driving EndMT in PAH/PH are still elusive. EndMT is induced by the activated or upregulated SNAI1/2/3, ZEB1/2, or TWIST1/2^1,7^. SNAI1, a transcription factor or transcription repressor, directly represses endothelial cell marker genes, such as CDH5 (VE-cadherin). Besides the initial transcription factors, molecular signatures of EndMT include downregulation of EC-specific markers along with upregulation of SMC-specific and fibroblast (FB)-specific markers. These molecular changes in LVEC result in actin filaments reorganization and switching from cobblestone to spindle-shape morphology. Additionally LVEC undergoing EndMT lose cell-cell junctions which allow stationary cells to migrate.

Inflammation is considered as an important trigger for the development of pulmonary vascular remodeling in PAH/PH through different mechanisms^8,9^. Published literature demonstrates that inflammatory cytokines, such as transforming growth factor-β (TGF-β), tumor necrosis factor α (TNF-α), and interleukin-1β (IL-1β), induce EndMT^7,10,11^. We recently demonstrated that TGF-β_1_ promoted EndMT in normal human LVEC to a level similarly seen in LVEC from patients with PAH^5^. Therefore, TGF-β-mediated EndMT may contribute to the development of vascular remodeling in PAH/PH lungs. It has been proposed that EndMT is involved in the complex interactions between inflammatory stress and endothelial dysfunction^10^.

Regulation of Ca^2+^ signaling is involved in EC function however the role of Ca^2+^ signaling in EndMT has not been fully studied^12,13^. In LVEC, Ca^2+^ homeostasis is maintained by two distinct Ca^2+^ influx mechanisms: voltage-dependent and voltage-independent pathways^14,15^. Voltage-dependent Ca^2+^ entry (VDCE) is induced by membrane depolarization while voltage-independent Ca^2+^ entry is induced by store-operated Ca^2+^ entry (SOCE) and receptor-operated Ca^2+^ entry (ROCE).. Voltage-independent Ca^2+^ signaling is initiated by the stimulation of G protein-coupled receptors (GPCR) or receptor tyrosine kinases (RTK) by their extracellular ligands. The phospholipase C (PLC) is activated and two important second messengers, diacylglycerol (DAG) and inositol 1,4,5-trisphosphate (IP_3_), are produced^16^. DAG then opens receptor-operated Ca2+ channels (ROCC) and increases [Ca^2+^]_cyt_ through ROCE. In turn, IP_3_ binds to the IP_3_ receptor (IP_3_R) on the endoplasmic reticulum (ER) membrane to release Ca^2+^ from the ER to the cytosol via IP_3_R. Depletion of intracellular stored Ca^2+^ in the ER initiates the conformational change of stromal interaction molecule (STIM), a Ca^2+^ sensor on the ER membrane, to oligomerize and translocate to the ER-plasma membrane junction (puncta) to recruit calcium release-activated calcium channel 1 (Orai1). The STIM-Orai1 complex forms the store-operated Ca^2+^ channel (SOCC) for SOCE. It has been shown that SOCE contributes to EC proliferation and migration^17,18^, however, the role of SOCE in EndMT is still unclear. In this study we aimed to explore the role of endothelial Ca^2+^ signaling in EndMT using *in vitro* approach.

## Methods and Materials

### Human cell culture

Normal human LVEC, pulmonary arterial SMC and FB were purchased from Lonza. LVEC were cultured in T25 flasks pre-coated with 0.1% gelatin in VascuLife® basal medium without antimicrobials and phenol red supplemented with VascuLife® VEGF-Mv LifeFactors kit (Lifeline Cell Technologies, Cat#LL-0005) while SMC were cultured in 100-mm Petri dishes in VascuLife® basal medium supplemented with VascuLife® SMC LifeFactors kit (Lifeline Cell Technologies, Cat#LL-0014). FibroLife® basal medium supplemented with FibroLife® S2 LifeFactors kit (Lifeline Cell Technologies, Cat#LL-0011) was used to culture FB in 100-mm Petri dishes. Cell passages between 3 and 6 were used in the present studies. All cells were maintained at 37°C in 5% CO_2_ humidified atmosphere. LVEC were treated with vehicle or TGF-β_1_ (10 ng/ml, R&D Systems, Cat# 240-B-010) for 7 days as described before^5^ unless otherwise indicated. The membrane impermeant EGTA (2 mM) was added to the culture medium (1.8 mM Ca^2+^) to reduce or chelate extracellular free Ca^2+^ from 1.8 mM to ∼535 nM (Ca^2+^-free medium). Low-Ca^2+^ medium (LCM) was prepared by serial dilutions from Ca^2+^-free medium based on the calculated [Ca^2+^]_cyt_. The membrane-permeable BAPTA-AM (1µM) was added to the culture medium to chelate intracellular Ca^2+^. The pH value was measured after addition of the chemicals and adjusted to 7.4. In some experiments cells were treated with or without Orai1 inhibitor, AnCoA4 (50 µM, Sigma, Cat#532999), in the presence of TGF-β_1_ (10 ng/ml) for 7 days. For morphometric analysis cells were imaged and manually outlined using ImageJ software (NIH) to quantify cell area and perimeter. Circularity was calculated as [(4π(cell area)/(cell perimeter)^2^]. Elongation index was calculated as [(cell perimeter)^2^/(4π(cell area))].

### [Ca^2+^]_cyt_ fluorescent imaging

The measurement of [Ca^2+^]_cyt_ was performed at room temperature (22-24°C) as previously described^19^. Briefly, vehicle- and TGF-β_1_-treated human LVEC were grown on 35-mm diameter round glass coverslips. Cells were incubated with 4 μM fura-2 acetoxymethyl ester (fura-2/AM, Invitrogen, Cat# M1291) in HEPES-buffered solution (137 mM NaCl, 5.9 mM KCl, 1.8 mM CaCl_2_, 1.2 mM MgCl_2_, 14 mM glucose, and 10 mM HEPES, pH 7.4) for 1 hour. The Ca^2+^-free solution was prepared by replacing 1.8 mM CaCl_2_ with equimolar MgCl_2_ and adding 2 mM EGTA to chelate residual Ca^2+^. Intracellular Ca^2+^ concentrations were expressed as 340/380 fluorescence ratio within an area of interest in a cell recorded every 2 seconds using an inverted fluorescent microscope (Eclipse Ti-E, Nikon) equipped with EM-CC camera (Evolve, Photometrics) and NIS Elements 3.2 software (Nikon).

### RT-PCR

Total RNA samples were extracted from cells using QIAzol reagent (Qiagen) as previously described^5^. cDNA samples generated using High Capacity cDNA Reverse Transcription kit (Applied Biosystems, Cat# 00654321) were used as templates in semi-quantitative end-point PCR with Taq Polymerase Master Mix (Roche, Cat# 04728858001) in a T100 Thermal Cycler (BioRad). PCR products were load onto 1.5% agarose gel in Sub-Cell GT Cell Tank (BioRad) and separated by electrophoresis at 120V. PCR products were visualized under UV light by Photo Imager system (VWR) and the intensity was quantified by ImageJ software (NIH). cDNA was quantitatively measured in triplicates by qPCR using iTaq Universal SYBR Green Supermix (Bio-Rad, Cat# 1725121) in CFX384 Touch Real-Time PCR Detection System (Bio-Rad). Human forward and reverse primers used in the study: SNAI1 (F–5’-GTTCTTCTGCGCTACTGCTG-3’, R–5’-TTAGGTCTCAAATGGAAGGTCGT-3’); SNAI2 (F-5’-GACACATTAGAACTCACACGGG-3’, R-5’-GACCGACGACACATCGTGTG-3’); STIM1 (F-5’-GCTCCTCTGGGGACTCCT-3’, R-5’-CAATTCGGCAAAACTCTGCT-3’), STIM2 (F-5’-CTTGCCTTCCCCTGATCCAGA-3’, R-5’-GAGCCCAAGGTGAATACATTG CT-3’), Orai1 (F-5’-ACCTCGGCTCTGCTCTCC-3’, R-5’-GATCATGAGCGCAAACAGG-3’), GAPDH (F-5’-GCACCGTCAAGGCTGAGAAC-3’, R-5’-TGACCGCAGAAGTGGTGG TA-3’). GAPDH was used as an internal control in all experiments. The 2^-ΔΔCt^ method was used to calculate the relative level of mRNA.

### Immunoblotting

Total protein samples were isolated from cells using RIPA buffer as previously published^5^. Briefly, protein samples were mixed with 6 × SDS-sample buffer (Boston BioProducts, USA) and denatured for 7 min at 100°C. Then samples were separated on Bolt™ 4-12% Bis-Tris Plus polyacrylamide gels (Invitrogen, Cat# 4561045EDU) by electrophoresis at 160V and transferred to 0.45 µm nitrocellulose membrane (Bio-Rad, Cat# 1620115) at 10V in Mini Gel Tank (Invitrogen). The membrane was blocked with 5% non-fat milk in 1× Tris-Buffered Saline with 0.1% Tween 20 (TBS-T) for 1 h at room temperature and then incubated overnight at +4°C with primary antibody: anti-Snai1 (Cat# 3879, Cell Signaling), anti-STIM1 (Cat# 4119, Prosci), anti-STIM2 (Cat# S8572, Sigma), anti-Orai1 (Cat# ACC-060, Alomone), or β-actin antibody (Cat# sc-47778, Santa Cruz Biotechnology) as a loading control. The membrane was washed with 1x TBS-T and then incubated with the secondary anti-mouse (Cat# 7076S, Cell Signaling) or anti-rabbit (Cat# 7074S, Cell Signaling) antibody in 5% non-fat milk for 1 h at room temperature. The membrane was subsequently developed using an enhanced chemiluminescence detection system (ThermoFisher Scientific, Cat# 34095). Band intensities on the membrane were quantified using ImageJ software (NIH).

### Cell counting

Cells were plated on 12-mm coverslips in 24-well plates at a density of 1×10^4^ cells. Automated cell counting was performed at days 1, 3, 5 and 7 using Countess II (ThermoFisher Scientific). Briefly, cells were detached and mixed with Trypan Blue before the loading into the counting slide. The cells were allowed to settle for 30 sec after loading the sample to help ensure a uniform focal plane and accurate counts. The counting slide was inserted and autofocus was initiated. Adjustments were applied for fine control of data selection and the viable cell concentration was collected from both sides of the counting slide.

### Cell transfection

Cells were transfected in Opti-MEM Reduced Serum Medium (Gibco, Cat# 51985091) with either control siRNA (Cat# sc-37007, Santa Cruz Biotechnology), STIM1 siRNA (Cat# sc-76589, Santa Cruz Biotechnology) or Orai1 siRNA (Cat# sc-76001, Santa Cruz Biotechnology) at the concentration of 60 pM for 6 h. Lipofectamine RNAiMAX (Invitrogen, Cat# 13778075) was used as transfection reagent. Then the transfection complex was washed out and cells were treated with vehicle or TGF-β_1_ (10 ng/ml) for 7 days as described above.

### Immunofluorescence

Cells plated on 2-chamber glass slides were transfected and/or treated as indicated above. Cells were fixed with 4% paraformaldehyde in PBS for 20 min at room temperature and followed by blocking the endogenous peroxidase with 3% H_2_O_2_ in 100% methanol for 10 min at room temperature. 1x TBS was used for washing. For blocking non-specific binding the samples were incubated in 10% goat serum in PBS for 1 h with shaking at room temperature followed by the incubation with primary antibodies: mouse anti-SNAI1 (Santa Cruz, Cat# sc-271977), rabbit anti-SNAI2 (Cell Signaling, Cat# 9585S), rabbit anti-ACTA2 (Cell Signaling, Cat# 19245S) or mouse anti-PECAM1 (Cell signaling, Cat# 3528S) antibody in 10% goat serum in PBS with shaking overnight at +4°C. Next day cells were incubated with Alexa Flour™ 488-conjugated or Alexa Flour™ 568-conjugated secondary antibody (Thermo Scientific) in 0.2% BSA in PBS for 1 h at room temperature. After wash-out coverslips were mounted on glass slides using mounting medium with DAPI (Vector Lab. Inc, Cat# H-1500-10). Images were acquired using Zeiss Axioobserver microscope with Apotome camera.

### Statistics

The data are expressed as means ± standard error (SE). Statistical analysis was performed using paired or unpaired Student’s *t*-test between two groups or ANOVA and appropriate post hoc tests among three or more groups. Differences were considered significant at *p*≤0.05.

## RESULTS

### TGF-β_1_ increases SNAI1/2 in LVEC

EndMT converts slowly growing LVEC into highly proliferative and migratory mesenchymal cells. We recently demonstrated EndMT and higher proliferative index in LVEC isolated from PAH patients than in normal LVEC^5^. By comparing the proliferation rate of normal human lung vascular EC, SMC and FB, we found that lung vascular EC grow significantly slower than SMC or FB (**Fig. 1A**). At day 7, the number of EC was 1.85-fold and 2.45-fold lower than that of SMC and FB, respectively (p<0.001) (**Fig. 1A**).

**Figure 1.**
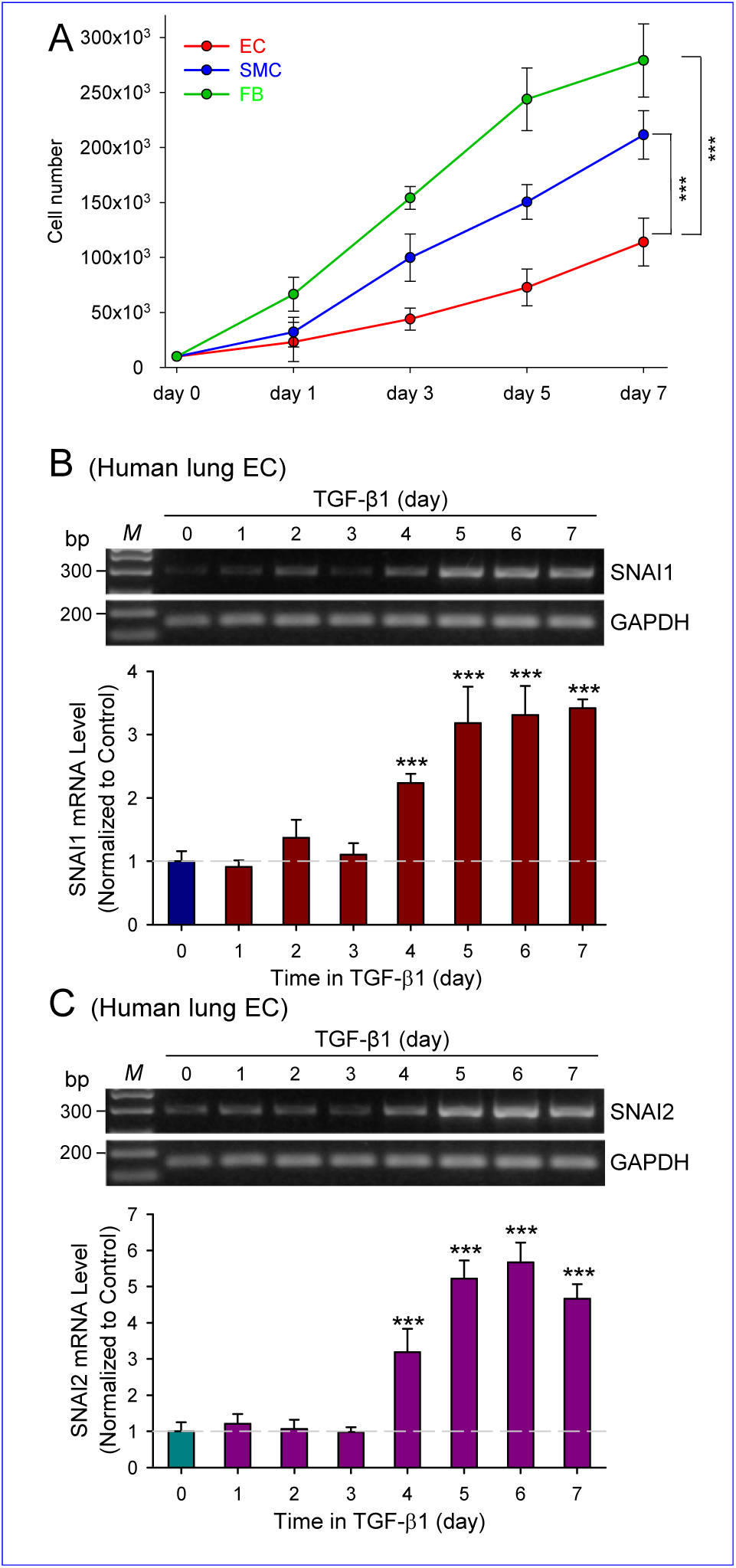
TGF-β_1_ increases SNAI1/2 in human LVEC in time-dependent manner. **A:** Cell numbers of normal human lung vascular EC, SMC and FB cultured for 1, 3, 5 and 7 days under the optimal conditions. **B, C:** Representative images (*top panel*) and summarized data (*bottom panel*) showing RT-PCR analysis of *SNAI1* (B) and *SNAI2* (C) time-course expression in LVEC treated with TGF-β_1_ (10ng/ml) for 7 days. Values are mean±SE (n=5 per group). \*\*\**p*<0.001. Statistical analysis was performed using One-Way ANOVA and post hoc test.

We previously reported that TGF-β_1_ treatment for 7 days induced EndMT due to upregulation of SNAI1/2 in normal human LVEC^5^. Using TGF-β_1_-induced EndMT as a model, we measured SNAI1/2 mRNA expression in cells treated with TGF-β_1_ every 24 h for 7 days. As shown in **Figure 1** (**B** and **C**), TGF-β_1_ increased SNAI1 (**Fig. 1B**) and SNAI2 (**Fig. 1C**) in a time-dependent manner. SNAI1 and SNAI2 levels at day 4 treatment with TGF-β_1_ were 2.23±0.15 and 3.19±0.65, respectively, which is 2-3 times higher than the level in non-treated cells (day 0; *p*<0.001). From day 5 to day 7 of TGF-β_1_ treatment, the mRNA levels of SNAI1 and SNAI2 further increased (from 2.23±0.15 and 3.19±0.65 to 3.42±0.14 and 4.67±0.40, respectively; *p*<0.001). These findings demonstrate that TGF-β_1_ upregulates SNAI as early as 4 days in human LVEC.

### Ca^2+^ is required for TGF-β_1_-mediated EndMT

The next set of experiments was designed to examine whether extracellular Ca^2+^ is required for TGF-β_1_-mediated upregulation of SNAI1/2. We treated LVEC with vehicle or TGF-β_1_ in the presence (2 mM EGTA, ∼535 nM Ca^2+^) or absence (0 mM EGTA, 1.8 mM Ca^2+^) of EGTA for the indicated period of time and then measured mRNA expression levels of SNAI1 and SNAI2. Our RT-PCR experiments showed that the long-term treatment of LVEC with TGF-β_1_ (7 days) resulted in a significant increase of SNAI1 and SNAI2 levels compared with control (**Fig. 2A**). Treatment of LVEC with vehicle in EGTA-containing medium for 7 days resulted in a dramatical decrease of SNAI1 and SNAI2 expression levels. The internal control (GAPDH) used for this experiment was also affected (data not shown). Short-term treatment of LVEC with TGF-β_1_ (for 4 hours) resulted in a significant increase of SNAI1 level, but not SNAI2 level, compared with the vehicle control (**Fig. 2B**). TGF-β_1_ was unable to increase SNAI1 in EGTA-containing medium. Since both SNAI1 and SNAI2 levels were significantly upregulated at day 4 in TGF-β_1_–treated cells (see Fig. 1B and C), we used 4 days as a time point to compare mRNA expression of SNAI/SNAI2 in LVEC treated with vehicle or TGF-β_1_ in the absence or presence of EGTA. As shown in **Figure 2** (**C-F**), both SNAI and SNAI2 were elevated after 4 days of TGF-β_1_ treatment, whereas chelation of extracellular free Ca^2+^ with EGTA abolished the TGF-β_1_ effect (**Fig. 2C** and **D**).

**Figure 2.**
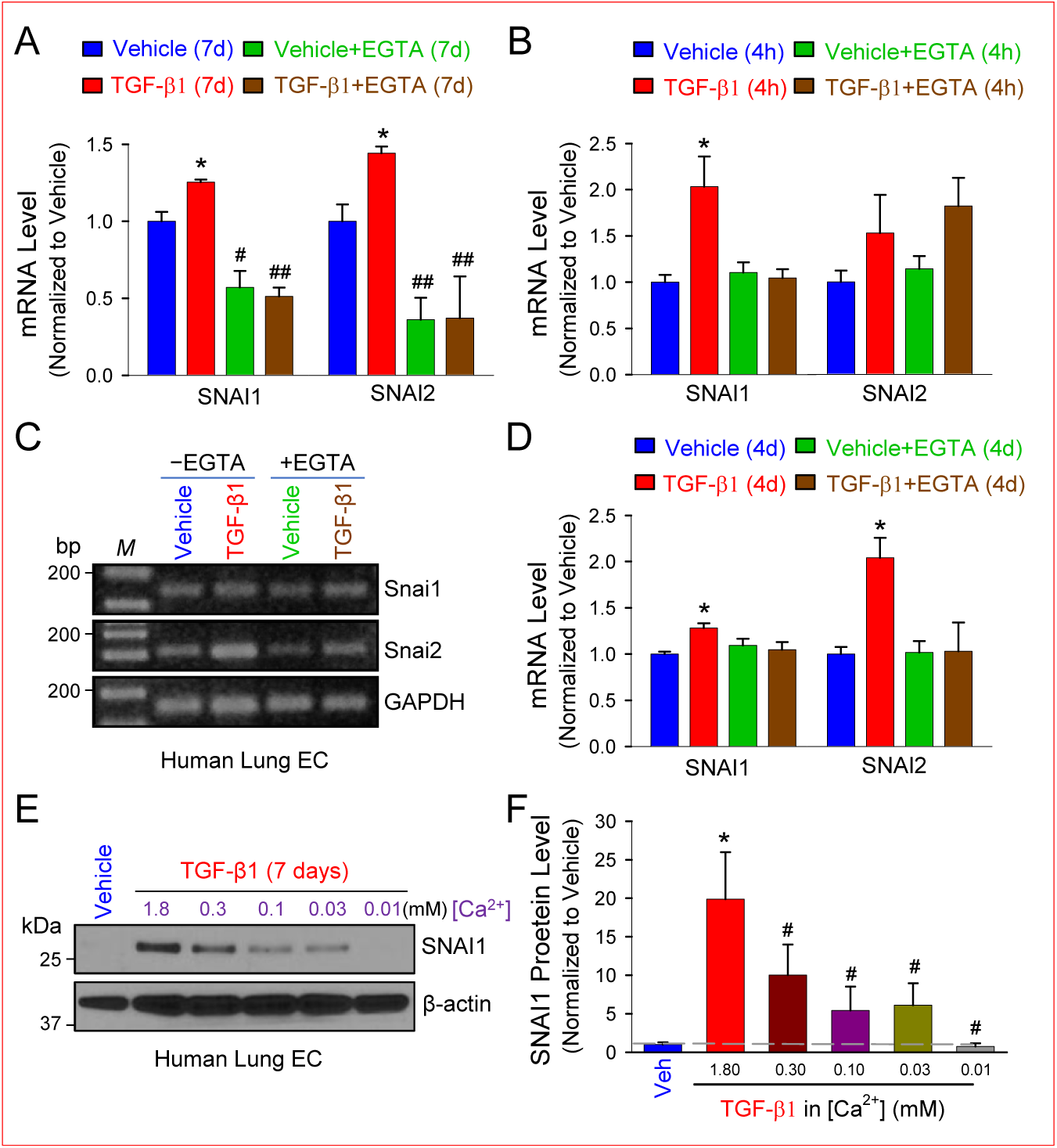
TGF-β_1_–mediated upregulation of SNAI depends on extracellular Ca^2+^. **A:** Real-time RT-PCR analysis of *SNAI1* and *SNAI2* expression (normalized to *GAPDH*) in LVEC treated with vehicle or TGF-β_1_ (10ng/ml) in the presence or absence of EGTA (2mM) for 7 days. **B:** Real-time RT-PCR analysis of *SNAI1* and *SNAI2* expression (normalized to *GAPDH*) in LVEC treated with vehicle or TGF-β_1_ (10ng/ml) in the presence or absence of EGTA (2mM) for 4 hours. **C, D:** Representative images (*left panel*) and summarized data (*right panel*) showing RT-PCR analysis of *SNAI1* and *SNAI2* expression (normalized to *GAPDH*) in LVEC treated with vehicle or TGF-β_1_ (10ng/ml) in the presence or absence of EGTA (2mM) for 4 days. **E, F:** Representative images (*left panel*) and summarized data (*right panel*) showing Western blot analysis of SNAI1 expression (normalized to β-actin) in LVEC treated with vehicle or TGF-β_1_ (10ng/ml) in the presence (0.03, 0.1, 0.3, or 1 mM EGTA; ∼0.3 mM, ∼0.1 mM, ∼0.03 mM or ∼0.01 mM Ca^2+^ respectively) or absence (0 mM EGTA, 1.8mM Ca^2+^) of EGTA for 7 days. Values are mean±SE. \**p*<0.05 vs Vehicle; ^#^*p*<0.05, ^##^*p*<0.01 vs TGF-β_1_. Statistical analysis was performed using One-Way ANOVA and post hoc test.

To further confirm that the upregulation of SNAI by long-term treatment of TGF-β_1_ is Ca^2+^ dependent, we treated LVEC with vehicle or TGF-β_1_ in the presence (0.03, 0.1, 0.3, or 1 mM EGTA, which correspond to ∼0.3 mM, ∼0.1 mM, ∼0.03 mM or ∼0.01 mM free [Ca^2+^], respectively) or absence of EGTA (0 mM EGTA, 1.8 mM Ca^2+^) for 7 days (**Fig. 2E**). Our Western blot experiments showed that SNAI1 protein level was markedly enhanced by TGF-β_1_ (to 19.86±6.11), but was significantly decreased (to 10.03±3.97) in cells incubated in 0.3-mM Ca^2+^ medium (**Fig. 2F**). The SNAI1 level in TGF-β_1_-treated cells was further decreased in EGTA-containing medium with approximately 0.01 mM Ca^2+^. Thus, chelation of extracellular free Ca^2+^ with EGTA diminished TGF-β_1_-induced increase in SNAI in a dose-dependent manner.

Additionally, we performed immunocytochemistry experiments to determine EndMT profile in LVEC treated with vehicle or TGF-β_1_ in the presence (1 mM EGTA; ∼0.01 mM Ca^2+^) or absence (0 mM EGTA, 1.8 mM Ca^2+^) of EGTA for 7 days. As shown in **Figure 3A**, SNAI1, SNAI2, and ACTA2 signals were not prominent while PECAM1 staining revealed an expected pattern at cell-cell borders in vehicle-treated cells. TGF-β_1_ increased SNAI1, SNAI2 and ACTA2, but reduced PECAM1 expression level (**Fig. 3A** and **B**). Chelation of extracellular free Ca^2+^ with cell-impermeable EGTA decreased SNAI1, SNAI2 and ACTA2 levels similar to the level in vehicle group, but had no effect on PECAM1, compared to the TGF-β_1_-treated group. Moreover, EGTA decreased cell numbers, but cells could tolerate the low extracellular Ca^2+^ concentration for the treatment duration (**Fig. 3A**). To examine the effect of intracellular Ca^2+^ or intracellularly-stored Ca^2+^ on TGF-β-mediated changes in SNAI1/2, ACTA2 and PECAM1, we also treated the cells with TGF-β_1_ for 7 days in the presence of the cell-permeable BAPTA-AM (1 µM), to chelate cytosolic free Ca^2+^ and intracellularly-stored Ca^2+20,21^. Treatment of the cells with the combination of TGF-β_1_ and BAPTA-AM (1 µM) for 7 days abolished TGF-β_1_-induced EndMT and significantly inhibited TGF-β_1_-induced increases in SNAI1/SNAI2 and ACTA2 (**Fig. 3A** and **B**). Importantly, BAPTA-AM partially rescued PECAM1 in TGF-β_1_-induced cells. It is worth to note that we observed the nuclear translocation of SNAI1 (as indicated by cyan color) suggesting its activation as well as SNAI1 and ACTA2 co-localization (as indicated by yellow color) upon TGF-β_1_ treatment (**Fig. 3C)**.

**Figure 3.**
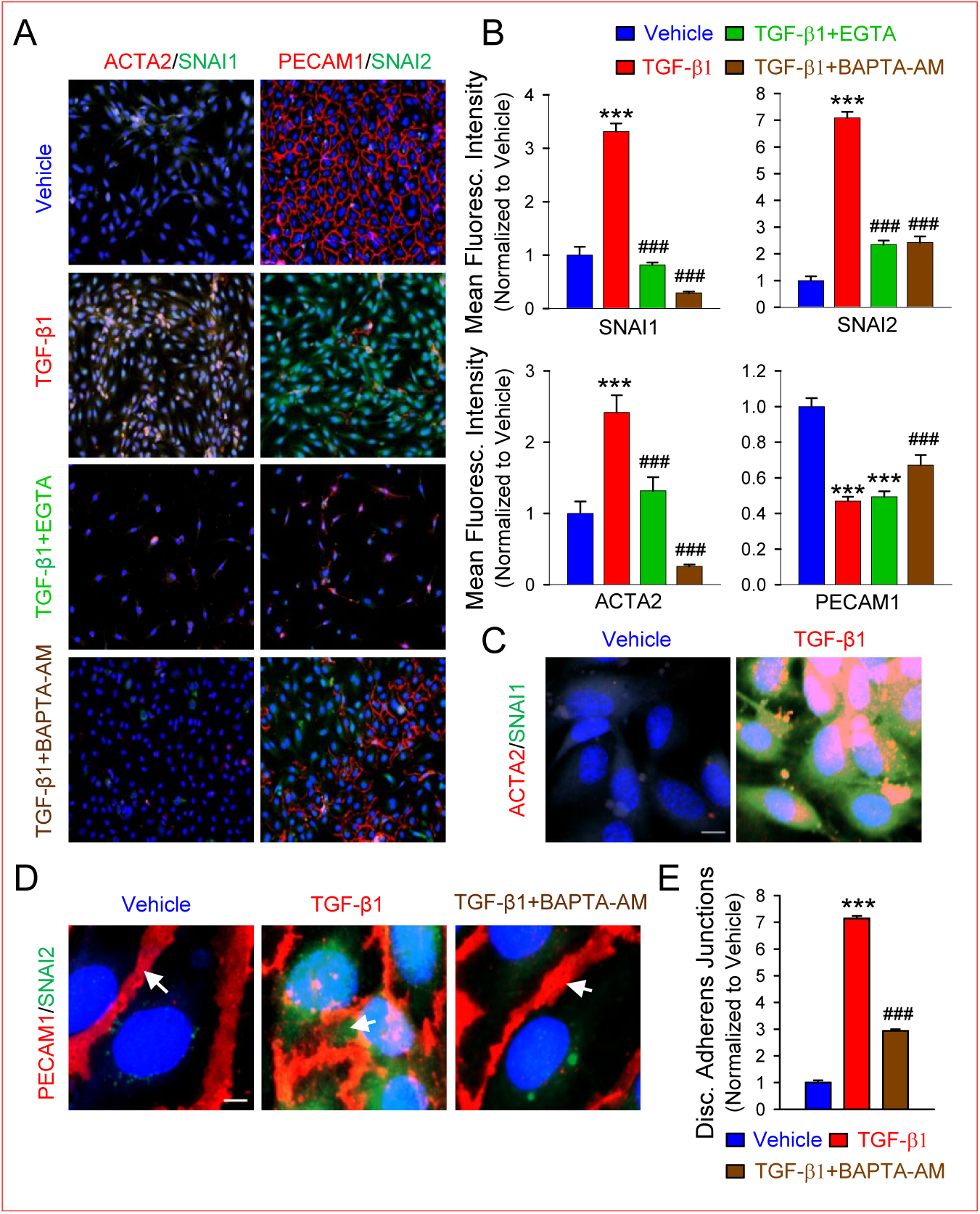
TGF-β_1_ requires extracellular and intracellular Ca^2+^ to induce EndMT. **A, B:** Representative images (A) and summarized data (B) showing mean fluorescence intensity in LVEC treated with vehicle or TGF-β_1_ (10ng/ml) in the presence (1mM EGTA, ∼0.01 mM Ca^2+^) or absence of EGTA (0 mM EGTA, 1.8mM Ca^2+^) or BAPTA-AM (1µM) for 7 days and stained for SNAI1, SNAI2, ACTA2 and PECAM1. **C:** Representative images showing mean fluorescence intensity in LVEC treated with vehicle or TGF-β_1_ (10ng/ml) for 7 days and stained for SNAI1 and ACTA2. Scale bar is 10µm. **D, E:** Representative images (D) and summarized data (E) showing mean fluorescence intensity in LVEC treated with vehicle or TGF-β_1_ (10ng/ml) in the presence or absence of BAPTA-AM (1µM) for 7 days and stained for PECAM1 and SNAI2. Scale bar is 5 µm. Values are mean±SE (n=6-12 per group). \*\*\**p*<0.001 vs Vehicle; ^###^*p*<0.001 vs TGF-β_1_. Statistical analysis was performed using One-Way ANOVA and post hoc test.

We previously reported that TGF-β_1_ dramatically disrupted the continuous adherens junctions in LVEC^5^. Next, we aimed to determine whether BAPTA-AM is able to attenuate TGF-β_1_ effect. In normal LVEC, PECAM1 localizes to adherens junctions that form continuous cell-cell contacts however LVEC in TGF-β_1_ group exhibited a significant increase in discontinuous adherens junctions (**Fig. 3D** and **E**). Moreover, we observed TGF-β_1_-mediated nuclear translocation of SNAI2 (as indicated by cyan color) which was suppressed by BAPTA-AM. These results lead us to conclude that Ca^2+^ influx and release are both required for or involved in TGF-β_1_-mediated SNAI upregulation and EndMT in human LVEC.

### TGF-β_1_ enhances SOCE and upregulates SOCC in normal human LVEC

To study Ca^2+^ regulating mechanisms responsible for TGF-β_1_-mediated EndMT, we measured [Ca^2+^]_cyt_ in control (non-treated), vehicle- or TGF-β_1_-treated cells (3 days). We used cyclopiazonic acid (CPA, 10 μM), a reversible inhibitor of the sarcoplasmic (SR) and endoplasmic reticulum (ER) Ca^2+^ pump (SERCA) that depletes Ca^2+^ from the intracellular stores. Extracellular application of CPA in the Ca^2+^-free solution (0Ca) resulted in a transient increase in [Ca^2+^]_cyt_ (1^st^ peak) due apparently to Ca^2+^ release or Ca^2+^ leakage from the intracellular Ca^2+^ stores (**Fig. 4A**). Restoration of extracellular Ca^2+^ (1.8Ca) caused a second increase in [Ca^2+^]_cyt_ (2^nd^ peak) which was due apparently to SOCE. As shown in **Figure 4B**, no significant changes in the [Ca^2+^]_cyt_ increases due to Ca^2+^ release/leakage (1^st^ peak) and to SOCE (2^nd^ peak) were found between cells treated with or without TGF-β_1_ for 3 days. However, the treatment with TGF-β_1_ for 7 days resulted in the enhancement of SOCE (2^nd^ peak) (**Fig. 4C**). CPA-mediated rise in [Ca^2+^]_cyt_ due to SOCE (2^nd^ peak) was significantly greater in TGF-β_1_-treated cells than in control cells (p<0.05) or vehicle-treated (p<0.05) cells **(Fig. 4D)**. To determine whether TGF-β_1_ affects SOCC, we performed a time-course of TGF-β_1_ treatment in LVEC and quantified the mRNA expression of STIM and Orai1 every 24 h for 7 days (**Fig. 4E)**. Our data showed that TGF-β_1_ upregulated STIM1, STIM2 and Orai1 in time-dependent manner (**Fig. 4F-H**). STIM1 and STIM2 were significantly increased by TGF-β_1_ within the first 24 h (p<0.001) while Orai1 was elevated at the day 2 (p<0.001). The expression of Orai2 was not changed within the time-course of TGF-β_1_ (data not shown). Among the three Orai homologs, Orai1 contributes the most to mediate SOCE^22^. The RT-PCR results were confirmed by Western blot analysis. Thus, the protein levels of Orai1, STIM1 and STIM2 were elevated significantly in LVEC at day 7 of TGF-β_1_ treatment compared with control or vehicle-treated cells (**Fig. 4I** and **J**). Taken together, the observations from these experiments indicate that TGF-β_1_ promotes SOCE due to at least the upregulated STIM1/2 and Orai1.

**Figure 4.**
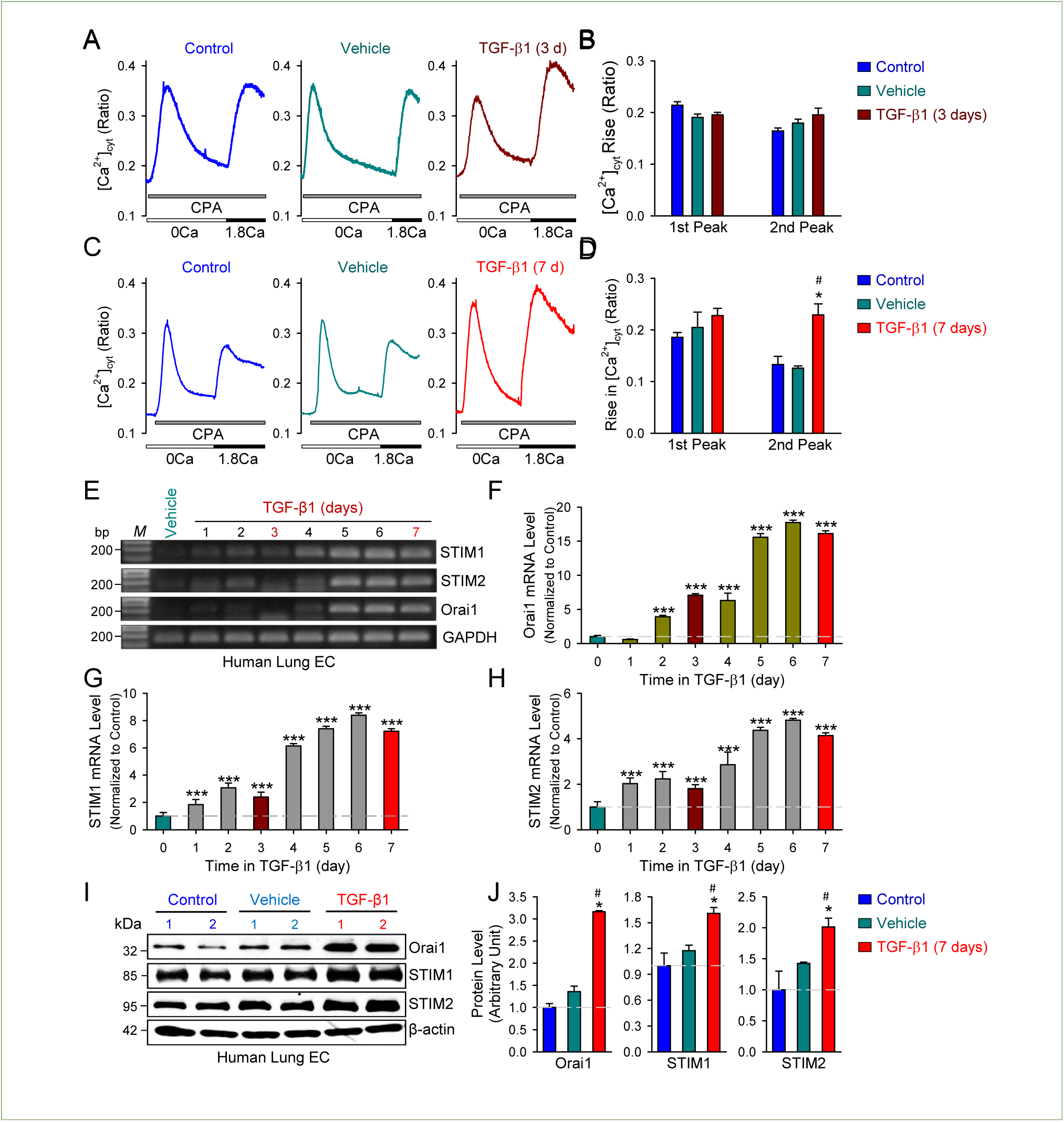
TGF-β_1_ enhances CPA-induced rise in [Ca^2+^]_cyt_ and upregulates STIM1/2 and Orai1. **A, B:** Representative tracings (A) and summarized data (B) showing changes in [Ca^2+^]_cyt_ before and during extracellular application of cyclopiazonic acid (CPA, 10 μM) in the absence (0Ca) or presence (1.8Ca) of 1.8 mM Ca^2+^ in non-treated LVEC (Control) or LVEC treated with vehicle or TGF-β_1_ (10ng/ml) for 3 days. **C, D:** Representative tracings (C) and summarized data (D) showing changes in [Ca^2+^]_cyt_ before and during extracellular application of CPA in the absence (0Ca) or presence (1.8Ca) of 1.8 mM Ca^2+^ in non-treated LVEC (Control) or LVEC treated with vehicle or TGF-β_1_ (10ng/ml) for 7 days (n=100-120 cells per group). **E-H:** Representative images (E) and summarized data (F-H) showing RT-PCR analysis of *Orai1, STIM1 and STIM2* time-course expression (normalized to *GAPDH*) in LVEC treated with vehicle or TGF-β_1_ (10ng/ml) for 7 days. **I, J:** Representative images (I) and summarized data (J) showing Western blot analysis of Orai1, STIM1 and STIM2 protein expression (normalized to β-actin) in non-treated LVEC (Control) or LVEC treated with vehicle or TGF-β_1_ (10ng/ml) for 7 days. Values are mean±SE (n=5-6 per group). \**p*<0.05, \*\*\**p*<0.001 vs Vehicle; ^#^*p*<0.05 vs Control. Statistical analysis was performed using One-Way ANOVA and post hoc test.

### SOCC knockdown abolishes TGF-β_1_-mediated EndMT

This set of experiments was designed to determine the role of SOCC in TGF-β_1_-induced EndMT. siRNAs specifically targeted STIM1 and Orai1 were used to transfect LVEC prior to the treatment with TGF-β_1_. As expected, control (transfected with Control siRNA and treated with Vehicle) cells exhibit little to no expression of SNAI1, SNAI2 and ACTA2, while PECAM1 was highly expressed at cell-cell borders (**Fig. 5A**). Upon TGF-β_1_ treatment for 7 days, cells underwent EndMT were determined by the significant upregulation of SNAI1, SNAI2 and ACTA2 (**Fig. 5B**). Moreover, PECAM1 level was significantly reduced by TGF-β_1_ (p<0.001). Either STIM1 or Orai1 knockdown was able to markedly decrease TGF-β_1_-mediated activation of SNAI1, SNAI2 and ACTA2. PECAM1 expression was significantly restored in both TGF-β_1_+Orai1 siRNA and TGF-β_1_+STIM1 siRNA groups compared with TGF-β_1_ group (**Fig. 5B**). Our Western blot experiments showed that knockdown efficiency was 57-72% in STIM1-tranfected cells (**Fig. 5C** and **D**) and 85-90% in Orai1-tranfected cells (**Fig. 5E** and **F**). TGF-β_1_ failed to upregulate SNAI1 expression in cells pre-transfected with either STIM1 or Orai1 siRNA. However, we did not observe any effect of Orai1 or STIM1 silencing on TGF-β_1_-induced discontinuous adherent junctions indicating that EndMT was not fully blocked upon STIM1 or Orai1 knockdown (**Fig. 5G** and **H**). Importantly, TGF-β_1_-induced SNAI2 nuclear translocation (as indicated by cyan color) was abolished in cells with STIM1 or Orai1 deficiency. These findings led us to conclude that endothelial STIM1 and Orai1 contribute to the development of TGF-β_1_-induced EndMT by forming SOC and enhancing Ca^2+^ influx via SOCE.

**Figure 5.**
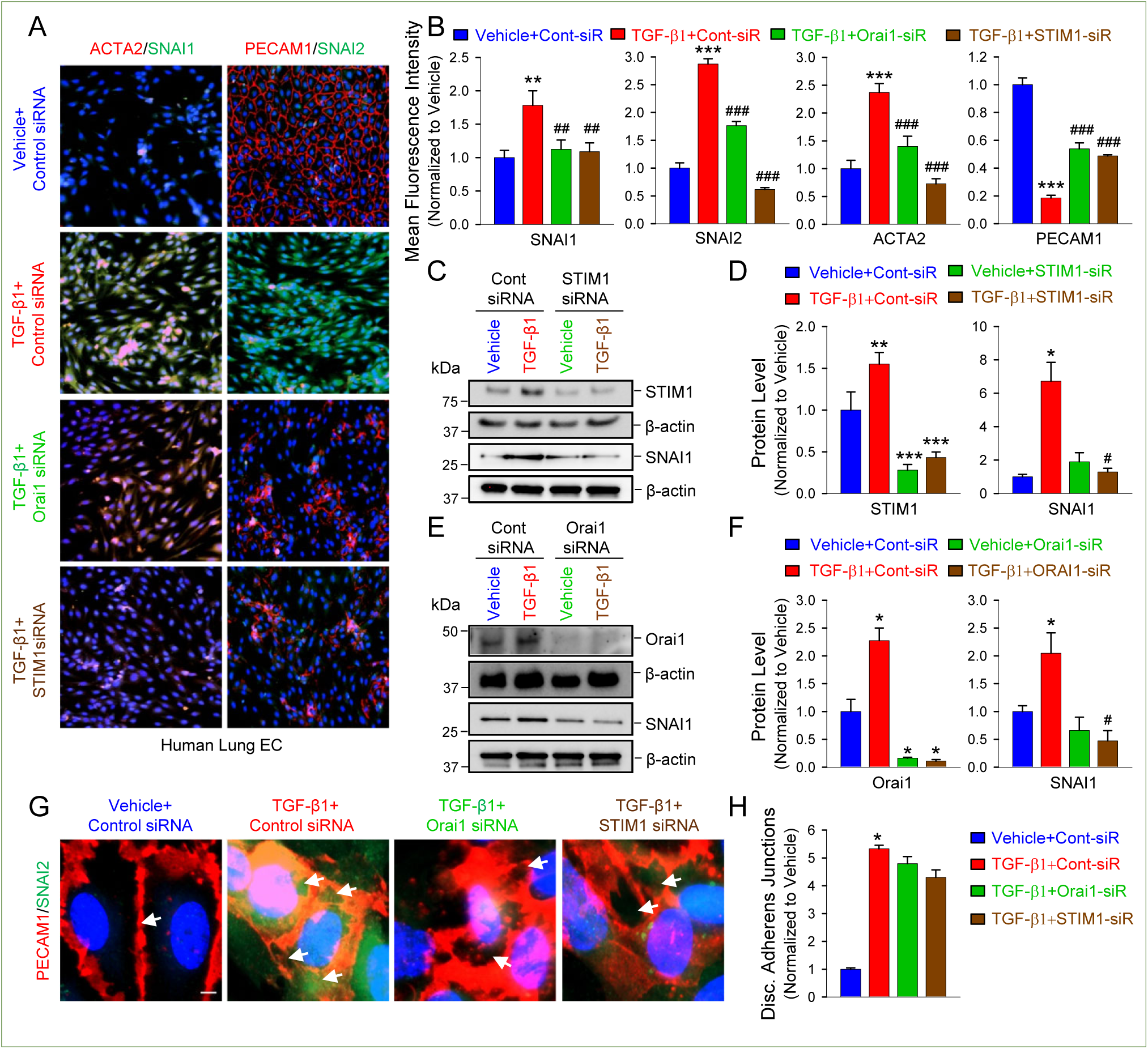
Orai1 or STIM1 knockdown inhibits EndMT but has no effect on the disrupted cell-cell contacts. **A, B:** Representative images (A) and summarized data (B) showing mean fluorescence intensity in LVEC transfected with control siRNA, Orai1 siRNA or STIM1 siRNA for 6 hours followed by the treatment with vehicle or TGF-β_1_ (10ng/ml) for 7 days and stained for SNAI1, SNAI2, ACTA2 and PECAM1 (n=6-12 per group). **C, D:** Representative images (C) and summarized data (D) showing Western blot analysis of STIM1 and SNAI1 protein expression (normalized to β-actin) in LVEC transfected with control siRNA or STIM1 siRNA for 6 hours followed by the treatment with vehicle or TGF-β_1_ (10ng/ml) for 7 days (n=5-6 per group). **E, F:** Representative images (E) and summarized data (F) showing Western blot analysis of Orai1 and SNAI1 protein expression (normalized to β-actin) in LVEC transfected with control siRNA or Orai1 siRNA for 6 hours followed by the treatment with vehicle or TGF-β_1_ (10ng/ml) for 7 days (n=5-6 per group). **G, H:** Representative images (G) and summarized data (H) showing mean fluorescence intensity in LVEC transfected with control siRNA, Orai1 siRNA or STIM1 siRNA for 6 hours followed by the treatment with vehicle or TGF-β_1_ (10ng/ml) for 7 days and stained for PECAM1 and SNAI2 (n=6-12 per group). Scale bar is 5µm. Values are mean±SE. \**p*<0.05, \*\**p*<0.01, \*\*\**p*<0.001 vs Vehicle+Control siRNA; ^#^*p*<0.05, ^##^*p*<0.01, ^###^*p*<0.001 vs TGF-β_1_+Control siRNA. Statistical analysis was performed using One-Way ANOVA and post hoc test.

### AnCoA4 inhibits TGF-β_1_-induced EndMT

Recently, AnCoA4 has been established as a SOCE-specific inhibitor that blocks STIM1-Orai1 interaction in pulmonary arterial endothelial cells^23^. Blockade of SOCC (STIM1/Orai1) or inhibition of SOCE with AnCoA4 (50 µM) attenuated TGF-β_1_-stimulated upregulation of SNAI1, SNAI2 and ACTA2 by 50.3%, 89.9% and 39.6%, respectively compared with TGF-β_1_ treatment alone (**Fig. 6A** and **B**). Importantly, AnCoA4 treatment not only significantly increased PECAM1 level by 123.1% compared with TGF-β_1_ treatment alone (**Fig. 6B**) but also restored continuous adherent junctions (**Fig. 6C** and **D**). Also AnCoA4 was able to abrogate SNAI2 nuclear translocation in TGF-β_1_-treated cells. Additionally, we evaluated AnCoA4 effect on TGF-β_1_-induced changes in cell morphology (**Fig. 6E**). Cell area and perimeter were used to determine cell circularity and elongation index. Circularity equals to 1 indicates the perfect circle while circularity less than 1 is defined as a deviation from the perfect circle (irregular shape). Thus, normal human LVEC have 315.9±35.14 µm^2^ area, 84.83±6.18 µm perimeter and circularity equals to 0.58±0.06 indicating a typical “cobblestone” shape (**Fig. 6F** and **G**). As expected, TGF-β_1_ induced spindle-shape pattern and a loss of cell polarity by significantly decreasing circularity and increasing cell elongation. AnCoA4 significantly decreased the cell area from 808.90±80.07 µm^2^ to 330.26±47.87 µm^2^ (p<0.001), cell perimeter from 161.45±6.13 µm to 84.26±5.96 µm (p<0.001), and circularity from 0.39±0.03 to 0.57±0.044 (p=0.025) in TGF-β_1_-treated cells. TGF-β_1_-induced cell elongation by 45.36% (compared with vehicle-treated cells) was abolished in AnCoA4 group (**Fig. 6E-G**). Overall, the current findings demonstrated that STIM1-Orai1 interaction is required for EndMT while AnCoA4 may have a therapeutic potential to inhibit EndMT and protect endothelial phenotype in PH.

**Figure 6.**
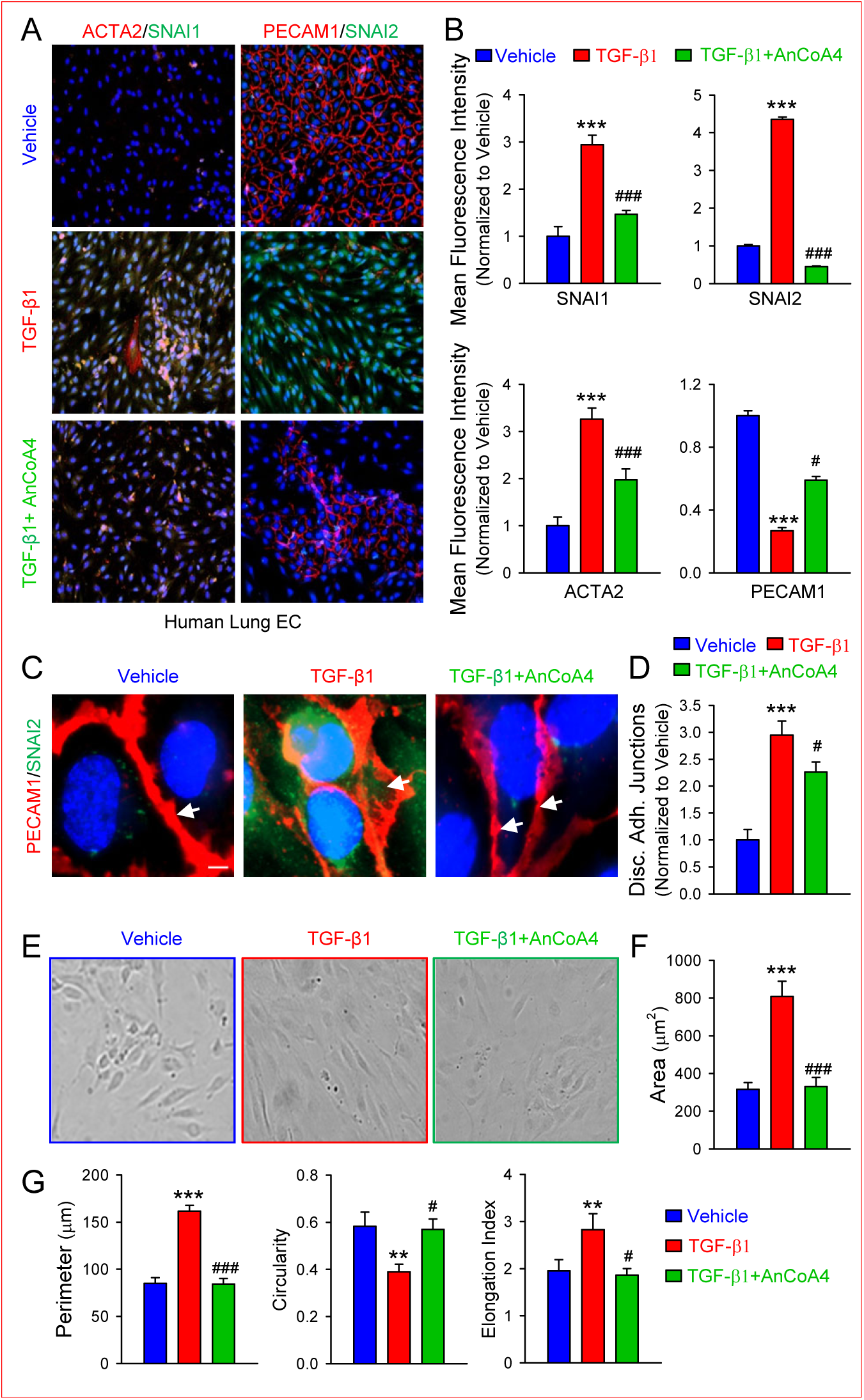
Orai1 inhibitor, AnCoA4, abolishes EndMT and restores endothelial phenotype. **A, B:** Representative images (A) and summarized data (B) showing mean fluorescence intensity in LVEC treated with vehicle or TGF-β_1_ (10ng/ml) in the presence of absence of AnCoA4 (50 µM) for 7 days and stained for SNAI1, SNAI2, ACTA2 and PECAM1. **C, D:** Representative images (C) and summarized data (D) showing mean fluorescence intensity in LVEC transfected with control siRNA, Orai1 siRNA or STIM1 siRNA for 6 hours followed by the treatment with vehicle or TGF-β_1_ (10ng/ml) for 7 days and stained for PECAM1 and SNAI2 (n=6-12 per group). Scale bar is 5µm. **E-G:** Representative images (E) and summarized data (F, G) showing morphological changes in cell area, perimeter, circularity and elongation index in LVEC treated with vehicle or TGF-β_1_ (10ng/ml) in the presence of absence of AnCoA4 (50 µM) for 7 days. Values are mean±SE (n=6-12 per group). \*\**p*<0.01, \*\*\**p*<0.001 vs Vehicle; ^#^*p*<0.05, ^###^*p*<0.001 vs TGF-β_1_. Statistical analysis was performed using One-Way ANOVA and post hoc test.

## DISCUSSION

The results from this study demonstrated that i) normal and fully differentiated EC grew significantly slower than SMC or FB within PA tissue; ii) TGF-β_1_, a well-known EndMT inducer, upregulated the transcription factor SNAI in a time-dependent manner; iii) TGF-β_1_-mediated increase in SNAI required extracellular and intracellular Ca^2+^ in lung vascular EC; iii) TGF-β_1_ enhanced SOCE-mediated rise in [Ca^2+^]_cyt_ by upregulating STIM1/2 and Orai1; iv) STIM1 or Orai1 knockdown significantly attenuated TGF-β_1_-mediated EndMT; v) blockade of STIM1-Orai1 interaction by AnCoA4 inhibited EndMT and restored EC-specific phenotype.

EndMT is a highly dynamic biological program activated in response to EC injury resulting in a phenotypic transition from fully differentiated EC to mesenchymal cells. Transitioning cells acquire the enhanced migratory abilities by losing endothelial cell-cell junctions which results in the increased migratory abilities. Additionally, LVEC undergoing EndMT program during the development and progression of PH could be a source for highly proliferative and apoptosis-resistant mesenchymal cells found in intimal and plexiform lesions from patients with PAH/PH. Our current study confirmed that EC had the slowest proliferation rate in comparison to other cell types within normal human PA (SMC or FB). LVEC isolated from patients or animals with PH are undergoing EndMT and converting into mesenchymal phenotype with the increased proliferative abilities^2,3,5^. Gaining higher proliferative ability allows mesenchymal cells to contribute to pulmonary vascular remodeling along with SMC and FB.

We used TGF-β_1_ to induce EndMT *in vitro* to determine the time-course of SNAI1/2 expression. In cancer studies TGF-β_1_ is well established stimulus for epithelial-to-mesenchymal transition (EMT), the process similar to EndMT^11,24^. Our highly sensitive qPCR experiments showed that SNAI1 upregulation by TGF-β_1_ occurs as early as 4 hours however SNAI2 level was not changed at this time-point. Less sensitive end-point PCR experiments demonstrated SNAI1/2 stimulation in TGF-β_1_ time-course at day 4. Zhou et al. reported that SNAI is highly unstable molecule with a short half-life time (25 min)^25^ which is consistent with our findings. In epithelial cells TGF-β_1_ was able to induce SNAI1 but not SNAI2 expression after 2 hours^26^ while stimulatory effect of TGF-β_1_ on SNAI1 expression in breast cancer cells occurred after 12-24 hours^27^. Molecular changes associated with EMT in these studies were observed at the time of SNAI activation. Our previous published studies documented EndMT in human LVEC treated with TGF-β_1_ for 7 days^5^. Taken together, the data indicate the cell specificity of TGF-β-induced changes in SNAI expression.

The role of Ca^2+^ signaling is widely studied in TGF-β_1_-induced EMT^28-30^ but this is, to the best of our knowledge, the first report about Ca^2+^ dependence of TGF-β_1_-induced EndMT. Here we provide a strong evidence that treatment of human LVEC with TGF-β_1_ enhanced [Ca^2+^]_cyt_ via SOCE. It has been well established that SOCE is important for TGF-β_1_-mediated EMT^26,27,30^. Thus, short-term TGF-β_1_ treatment for 24 hours promoted the 2^nd^ peak (SOCE) but not the 1^st^ peak (Ca^2+^ release) in colorectal and breast cancer cells^29,31^. In normal epithelial cells TGF-β_1_ had positive effect on [Ca^2+^]_cyt_ as early as 1 hour due to a rise in SOCE^32^. In that study stimulation of Ca^2+^ release due to chelation of extracellular Ca^2+^ by thapsigargin (SERCA pump blocker with higher potency than CPA^33,34^) did not show any differences between control and TGF-β_1_–treated cells. In electrophysiological experiments the authors demonstrated a significant increase in SOCC activity in cells exposed to TGF-β_1_^32^. Additionally, cytoskeleton dynamics in murine kidney cells treated with TGF-β_1_ was highly Ca^2+^-dependent^35^. This effect appears to be specifically mediated by SOCE since Ca^2+^ release from intracellular stores due to administration of thapsigargin did not affect actin cytoskeleton dynamics. These findings indicate that TGF-β_1_ affects SOCE and enhances [Ca^2+^]_cyt_ to reorganize actin filaments resulting in switching cell morphology towards mesenchymal spindle-shape phenotype which is in line with our results.

Here we provide compelling evidence that EndMT is a Ca^2+^-dependent event. The very low [Ca^2+^]_cyt_ in the culture medium containing 2 mM EGTA (∼535 nM Ca^2+^) could explain the inability of TGF-β-mediated SNAI increase in the current study. Furthermore, SNAI1 upregulation by TGF-β_1_ is dose-dependent on extracellular Ca^2+^. In our experiments TGF-β_1_ requires at least 30 µM extracellular Ca^2+^ to increase SNAI1 level. Interesting that other growth factors, including EGF and IGF^28,36^, were unable to induce EMT after chelation of intracellular Ca^2+^ with cell-permeable EGTA-AM or BAPTA-AM. The involvement of SOCE has been proposed in the development of EGF-induced EMT^37^.

In our study we determined the important role of extracellular and intracellular Ca^2+^ in the development of EndMT in lung EC. Both Ca^2+^ chelators attenuated TGF-β-induced increase in EndMT transcription factor SNAI and SMC-specific marker ACTA2. In contrast to EGTA, BAPTA-AM treatment was not stressful for cells and improved cell-cell contacts. Small number of cells was able to survive in the low extracellular Ca^2+^ environment (in EGTA-containing medium) with lost cell connections. We previously reported in human pulmonary arterial SMC and lung epithelial cells that short-term treatment with BAPTA (25 μM) significantly attenuated the increase in [Ca^2+^]_cyt_ induced by CPA by 33% while BAPTA-AM completely blocked CPA-induced rise in [Ca^2+^]_cyt_^19,38^. Recently we showed that BAPTA-AM (1 µM) was enough to chelate intracellular Ca^2+^ level in human lung vascular EC^39^ providing the dose justification for the current study. Zhou et al. observed the suppressed [Ca^2+^]_cyt_ in bone marrow macrophages treated with BAPTA-AM (0.5-2 µM) for 7 days^40^. All together the data indicate the critical role of endothelial Ca^2+^ signaling mechanisms in the initiation and progression of EndMT.

We were able to identify the key SOCC (STIM and Orai1)^41,42^ responsible for positive TGF-β_1_ effect on [Ca^2+^]_cyt_ in human lung EC. Similar to our study, TGF-β_1_ increased Orai1 and STIM1 protein expression within the first 24-36 h in colorectal and breast cancer cells^29,31^. However, in contrast to our results, siRNA against Orai1 did not prevent TGF-β_1_-induced upregulation of Snai1 in murine mammary epithelial cells^26^. Instead, Orai3 and STIM1 were responsible for Ca^2+^ influx during EMT initiation upon TGF-β_1_ treatment in those cells. Separate studies also confirmed that STIM1 silencing reduced TGF-β_1_ effect on SOCE, EMT molecular signature, migration, and calpain activity^31,32^. In breast cancer cells both STIM1 and STIM2 mediate SOCE to promote TGF-β_1_-induced EMT while STIM1 had greater effect than STIM2^27^. Nevertheless, STIM2 has been proposed to play an important role in non-SOCE (probably ROCE) following TGF-β_1_–mediated EMT. Although STIM1/2 and Orai1 upregulation in response to TGF-β_1_ occurs within the first 24-48 hours in the current study, we did not detect any changes in SOCE after 3 days of TGF-β_1_ treatment. We speculate that TGF-β_1_ effect depends on the formation of SOCC and Ca^2+^ influx via SOCE rather than on enhanced expression of STIM/Orai although we cannot rule out the role of other Ca^2+^ regulating mechanisms (i.e., ROCE, TRP channels) in EndMT program. These observations led us to conclude that SOCE plays a pivotal role in TGF-β_1_–mediated EndMT similar to that seen in EMT but the molecular mechanisms are likely cell-specific.

Pharmacological inhibition of SOCE with 2-APB or YM58483 resulted in dramatic reduction of [Ca^2+^]_cyt_ and EMT in TGF-β_1_-treated cells^27,29,31^. 2-APB is considered as a non-specific SOCE inhibitor while YM58483 is used as a specific Orai1/SOCE inhibitor. We introduced a novel specific Orai1 inhibitor AnCoA4 as a potential drug to inhibit EndMT and treat PH. Similar to our results with AnCoA4, YM58483 abolished LPS-induced endothelial injury and reduction in the expression of junctional proteins (i.e., VE-cadherin and occludin)^43^. In addition, YM58483 was able to reduce fibrosis in vitro and in vivo^44,45^. Orai1 inhibitors appear to have therapeutic potential to treat PH but the effect has been mostly studied in SMC^46-48^. Recently Li et al. reported the functional role of Orai1 in endothelium-dependent contraction during systemic hypertension^49^. Moreover, Orai channels have been linked to the increased endothelial permeability^50^. Mechanistically, endothelial Orai1 and/or SOCE could be involved in canonical or non-canonical TGF-β_1_ pathways. Thus, BAPTA-AM suppressed thapsigargin-stimulated EMT by modulating Smad2/3 signaling in lung cancer cells^30^. Also EMT seems to have a positive correlation with Ca^2+^-sensitive PI3K/AKT signaling (non-canonical TGF-β_1_ pathway)^26,36^. However, the further mechanistic studies are needed to define Ca^2+^ regulating signaling in EndMT during PH.

In conclusion, we propose that EndMT is dependent on Ca^2+^ signaling to convert LVEC into mesenchymal cells and contribute to pulmonary vascular remodeling in the development of PH. Moreover, STIM1/Orai1 interaction and Ca^2+^ influx via SOCE play a central role in the induction of EndMT, at least, in response to TGF-β_1_. Therefore, targeting endothelial Ca^2+^ pathways that regulate EndMT program could offer novel remodeling-focused therapeutic approaches to treat PH.

## DATA AVAILABILITY

The data presented in this study are available upon request from the corresponding author.

## AUTHOR CONTRIBUTIONS

A.B. and J.X.-J.Y. conceived and designed research; A.B., I.E., S.S., R.P., P.P.J., A.I., J.C. performed experiments; A.B., I.E., S.S. M.T., R.P., P.P.J., A.I., J.C. analyzed data; A.B., I.E., S.S., M.T., R.P., P.P.J., A.I., J.C., L.Y., S.P., C.P., W.T.W., Y.C., T.W., M.A., Y.S.P., C.M.P., J.X.-J.Y. interpreted results of experiments; A.B. and J.X.-J.Y. prepared figures, A.B. drafted manuscript, A.B., I.E., S.S., M.T., R.P., P.P.J., A.I., J.C., L.Y., S.P., C.P., W.T.W., Y.C., T.W., M.A., J.Y.-J.S., P.A.T., J.W., A.M., Y.S.P., C.M.P., J.X.-J.Y. edited and revised manuscript; A.B., I.E., S.S., M.T., R.P., P.P.J., A.I., J.C., L.Y., S.P., C.P., W.T.W., Y.C., T.W., M.A., J.Y.-J.S., P.A.T., J.W., A.M., Y.S.P., C.M.P., J.X.-J.Y. approved final version of manuscript.

## AUTHOR NOTES

- A. Babicheva and J.X.-J. Yuan contributed equally as corresponding and senior authors.
- Correspondence: A. Babicheva or J.X.-J. Yuan.

